# Decoding Object Weight from Electromyography during Human Grasping

**DOI:** 10.1101/2021.03.26.437230

**Authors:** Elnaz Lashgari, Atabak Pouya, Uri Maoz

**Affiliations:** Schmid College of Science and Technology, Chapman University, Orange, CA, USA; Crean College of Health and Behavioral Sciences; Institute for Interdisciplinary Brain and Behavioral Sciences Chapman University; Biology and Bioengineering California Institute of Technology; Anderson School of Management University of California Los Angeles

**Keywords:** Electromyography, Time-Domain features, PCA, Machine Learning algorithms, Pattern Recognition

## Abstract

Human urges, desires, and intentions manifest themselves in voluntary action. The final stages of such voluntary action are the muscle contractions that bring it about. Electromyography (EMG) signals measure such muscle contractions. Decoding action contents from EMG require advanced methods for detection, decomposition, processing, and classification and remains a challenge in neuroscience. This study presents a new, time-domain method of classifying EMG for grasping different types of objects. Our proposed method can classify objects with different weights with an accuracy of up to 90%. This progress in neuroscience affects other fields like physiology, brain-computer interfaces, robotics, and so on.

## 1. Introduction

Decoding grasp information from electromyography (EMG) is an ongoing challenge in neuroscience [1, 2], physiology and medicine [3], biomechanics [4], and brain-computer interfaces [5]. This also has applications in robotics [6, 7]. EMG signals can potentially enable seamless human-robot interaction. It is therefore of value to investigate hand posture synergies and goal-directed kinematic patterns of covariation observed between human hand joints. Dimensionality reduction is an important underlying concept for general geometrical interpretation of synergies. In other words, the number of degrees of freedom (DOF) of the human hand that can be controlled in an independent manner is smaller than the total number of available DOFs [8]. Therefore, finding an effective way of decoding the neuromuscular activities for accurate modeling and recognition of motion patterns will help convert human kinematic variables to smooth robotic motion. Understanding more about EMG could also benefit rehabilitation of stroke and other patients.

EMG pattern-recognition is a signal processing technology that has been proposed for object classification and for activity recognition. Previous studies have evaluated the ability of various EMG features and classifiers to recognize different features of the object [9–11]. Much of the research on the dynamics of grasping has focused on tracking frequency-domain changes in EMG using short-time Fourier transform and wavelets [12, 13].

In 2018, the authors mainly attempted binary classification for the two a priori most distinct classes by considering electroencephalography (EEG) and EMG signals and extracting frequency domain features with the same accuracy in our study [11]. Here, we used time-domain features and tested the extent to which machine-learning algorithms could classify the weight of a grasped object. Such classification is important for haptics, motion/path planning, robotic learning and training, rehabilitation, and investigation of human behavior. Further, we derive the classification accuracy of EMG from various muscles and compare them to the combination of all muscles. Our method can reliably classify objects of three different weights (165, 330, or 660 g) with high accuracy—over 90%. This result demonstrates the feasibility of multi-class classification of objects that differ only slightly and only with respect to weight.

## 2. Method

### 2.1 Data

In this study, we used the WAY_EEG_GAL dataset, which is freely available and is commonly used to test techniques to decode sensation, intention, and action from EMG and scalp EEG in humans performing a grasp-and-lift task [14]. In this paper, we focus on EMG data.

Twelve Participants performed a series of lifting movements in which the object’s weight (165, 330, or 660 g), surface friction (sandpaper, suede, or silk surface), or both varied. The EMG signals were sampled at 4 kHz. In each trial the participant was cued to reach for the object, grasp it with the thumb and index finger, lift it straight up in the air and hold it for a couple of seconds, then put it back on the support surface, release it, and, lastly, return the hand to a designated rest position (Figure 1-a). Four sensors were used to record kinematics and forces: F1-F2 correspond to force/torque sensors (ATI Nano), with Fx corresponding to lift force and Fz to grip force (Figure 1-b).

**Figure 1:**
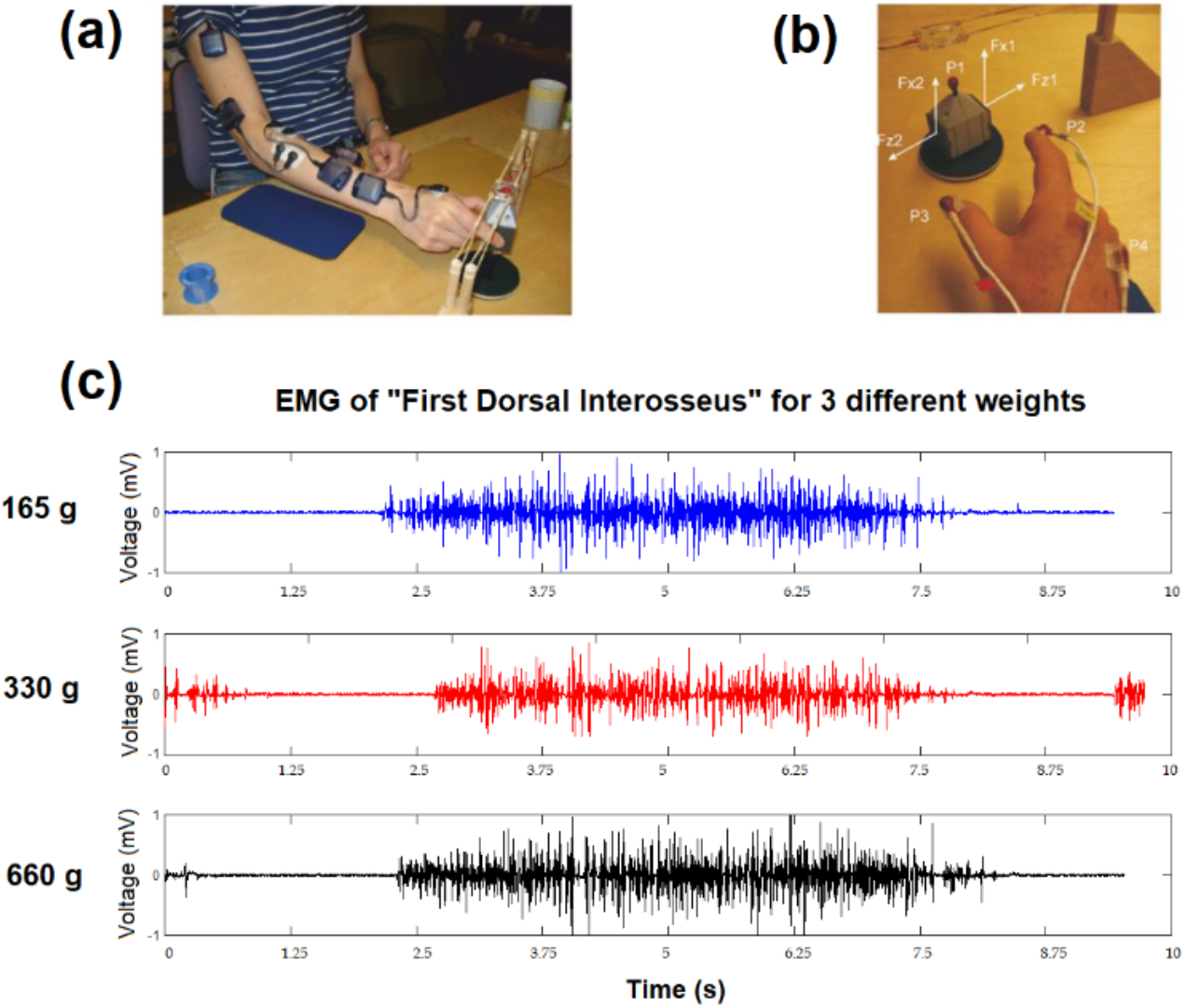
(a) Steps of the experiment: rest position, grasping, holding and releasing. (b) Force and position sensors (c) Raw EMG signals of First Dorsal Interosseous muscle for three different weights (165 g, 330 g, and 660 g)

We considered all available 2,204 trials of EMG signals, containing trials with different weights (700 trials for 165 g, 935 trials for 330 g, and 569 trials for 660 g) and constant sandpaper surface friction. Five EMG electrodes recorded the activity from 5 muscles (Anterior Deltoid, Brachioradialis, Flexor Digitorum, Common Extensor Digitorum, and First Dorsal Interosseous). The experiment contained other sessions, with different surface materials. We did not use those sessions. The raw EMG signals for the first dorsal interosseous for 3 different weights for Subject 1 are shown in Figure 1-c. All following single-subject data are from Subject 1. The signals were contaminated by various types of noise and artifacts. Therefore, pre-processing was necessary and important before feature extraction, especially for classes that are not highly distinct from one another. A recent publication classified the two a priori most distinct classes (165 and 660 g) by considering frequency features [11].

### 2.2 Pre-processing

It is difficult to obtain high-quality electrical signals from EMG because the signals typically have low amplitude (in the range of mV) and are easily corrupted by noise during recording. The EMG signal should, therefore, be pre-processed to suppress the noise before feature extraction. The most conventional technique for de-noising are filtering or smoothing. Filtering EMG data helps to remove high-frequency artifacts and low-frequency drifts. We thus used a Butterworth (4^th^ order) high-pass filter with a cut-off frequency of 450 Hz, followed by a low-pass Butterworth (4^th^ order) filter with a cut-off frequency of 5 Hz, with zero-phase on the full-wave rectified signals from each muscle. Finally, we normalized the signal by taking the z-score of each sample across trials (Figure 2).

**Figure 2:**
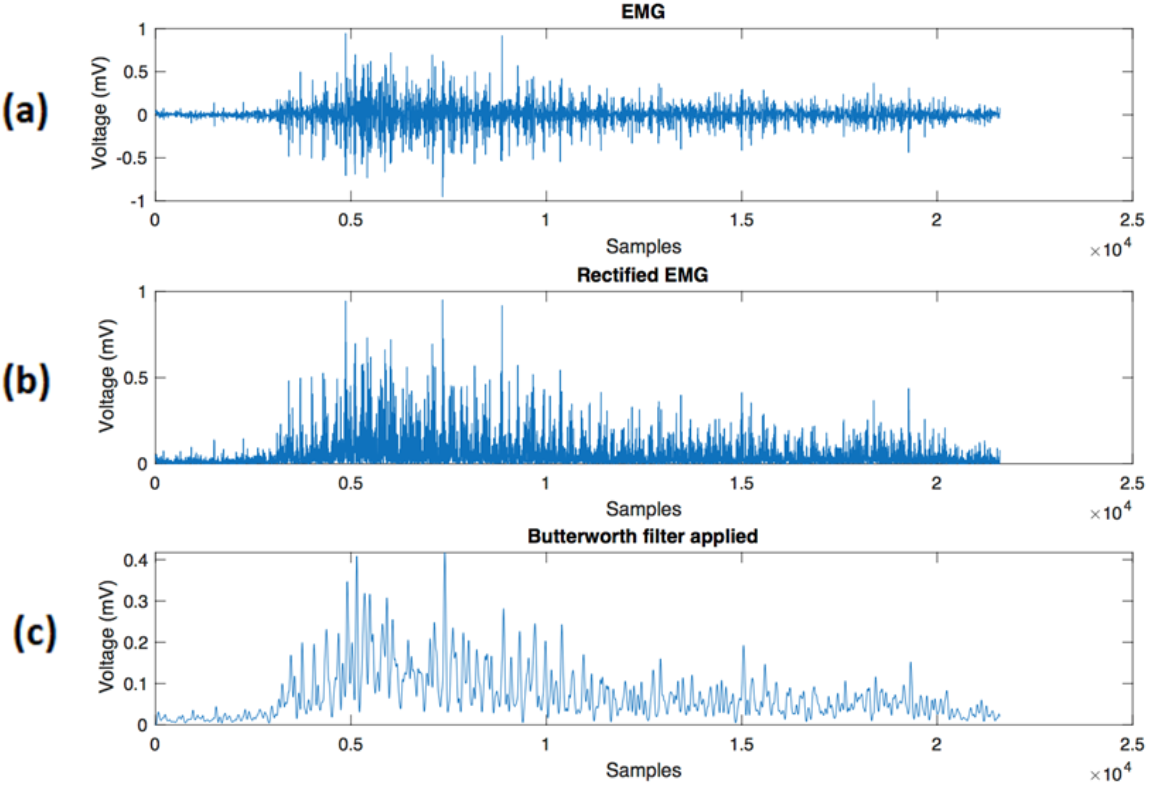
Pre-processing steps for an EMG signal from the First Dorsal Interosseous muscle: (a) Raw signal (b) Rectified signal (c) Signal after Butterworth (4th order) high-pass filter with cut-off frequency 450 Hz, followed by a 4^th^ order Butterworth low-pass filter with cut-off frequency 5 Hz with zero-phase.

### 2.3 Data selection

The original dataset is in MAT format (Mathworks MATLAB data files). Each trial took approximately 10 seconds to complete. We found no informative data before movement onset. To find the best informative segment, we locked on to the moment of touch (Figure 3) and picked the window from this point to 4 seconds after that (Figure 3). To identify the moment of touch, we used the moment when the normal force had increased above 4 times the standard deviation of the normal force during hand movements [14]. In the next step, we used a sliding window to find the most informative part of this 4 seconds segment.

**Figure 3:**
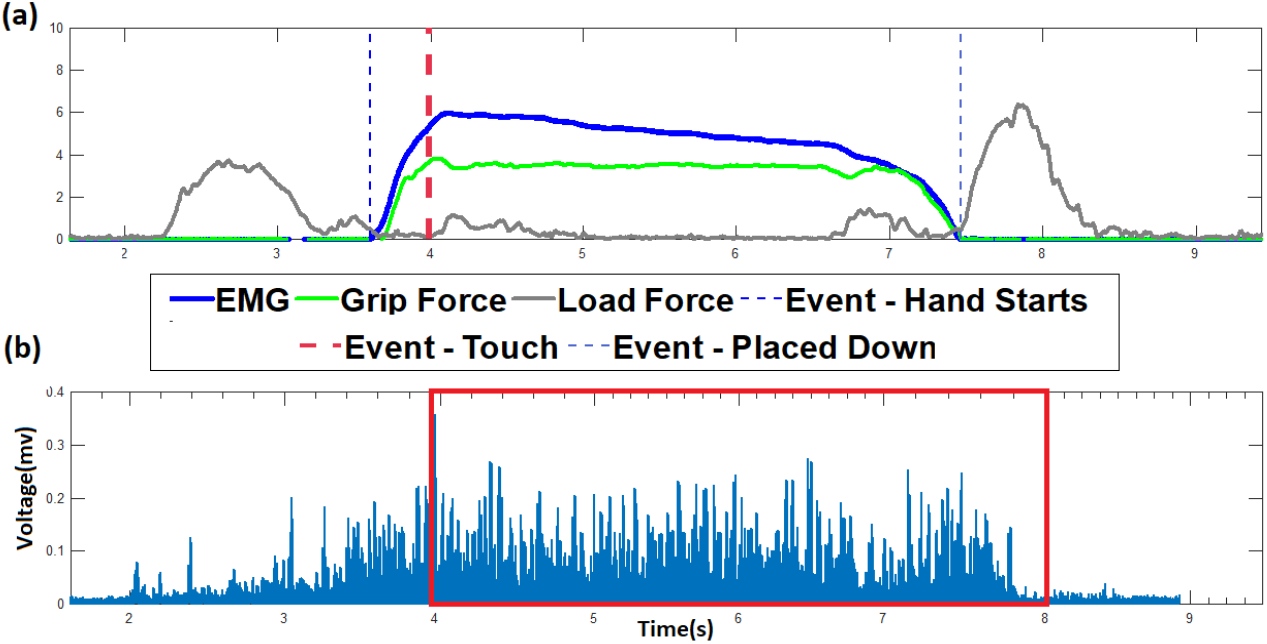
An example of EMG segmentation for the Brachioradialis muscle of Subject 1, based on the touch event. a) Grip Force, Load Force, and Event-touch plot. b) Raw EMG signal of Brachioradialis. The red rectangle shows the 4 seconds EMG signal after touch event point.

### 2.4 Feature extraction

In this study, we consider the time-domain features. Time-domain features or linear techniques are the most popular in EMG pattern recognition because of their computational simplicity [15–17]. Each sample of our trial includes the amplitude of EMG signal of the 5 muscles (see Data subsection of Methods). We consider the amplitude of each sample as a feature. Our results suggest that the Brachioradialis had the highest classification accuracy for detecting different weight classes. Also, the results reveal that considering the combination of all 5 muscles leads to better accuracy (Table 1).

**TABLE I.**
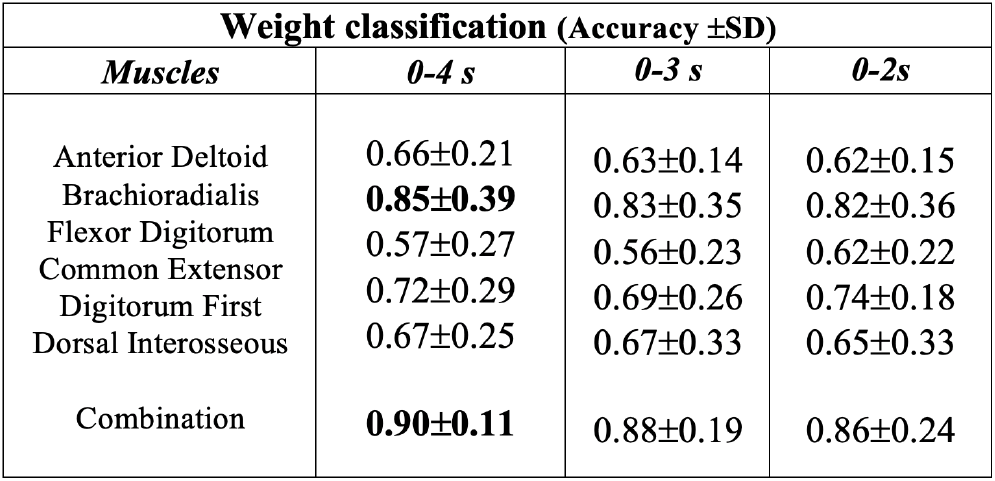
The accuracy of linear SVM for three classes of different muscles

### 2.5 Dimensionality reduction

Each 4 seconds segment consists 16,000 samples. We consider the amplitude of the samples as a feature set. As the number of features of a data increases, it becomes more difficult to process and analyze it. Dimension reduction is a solution to the curse of dimensionality when dealing with vector data in high-dimensional spaces. In this study, Principal Component Analysis (PCA) is used to reduce the dimension of the data by finding the minimum set of largest principal subspace that accounts for at least some pre-defined variance threshold [18–20].

The PCA algorithm results in a representation of the original data to a new coordinate system of lower dimension, using the first eigenvectors of the original data as axes. In our dataset 170 components were capable of describing 95% of the total variance. Figure 4 shows the first two prominent components of PCA. Instead of 16,000 features, these 170 components will be considered for the classification and further analysis.

**Figure 4:**
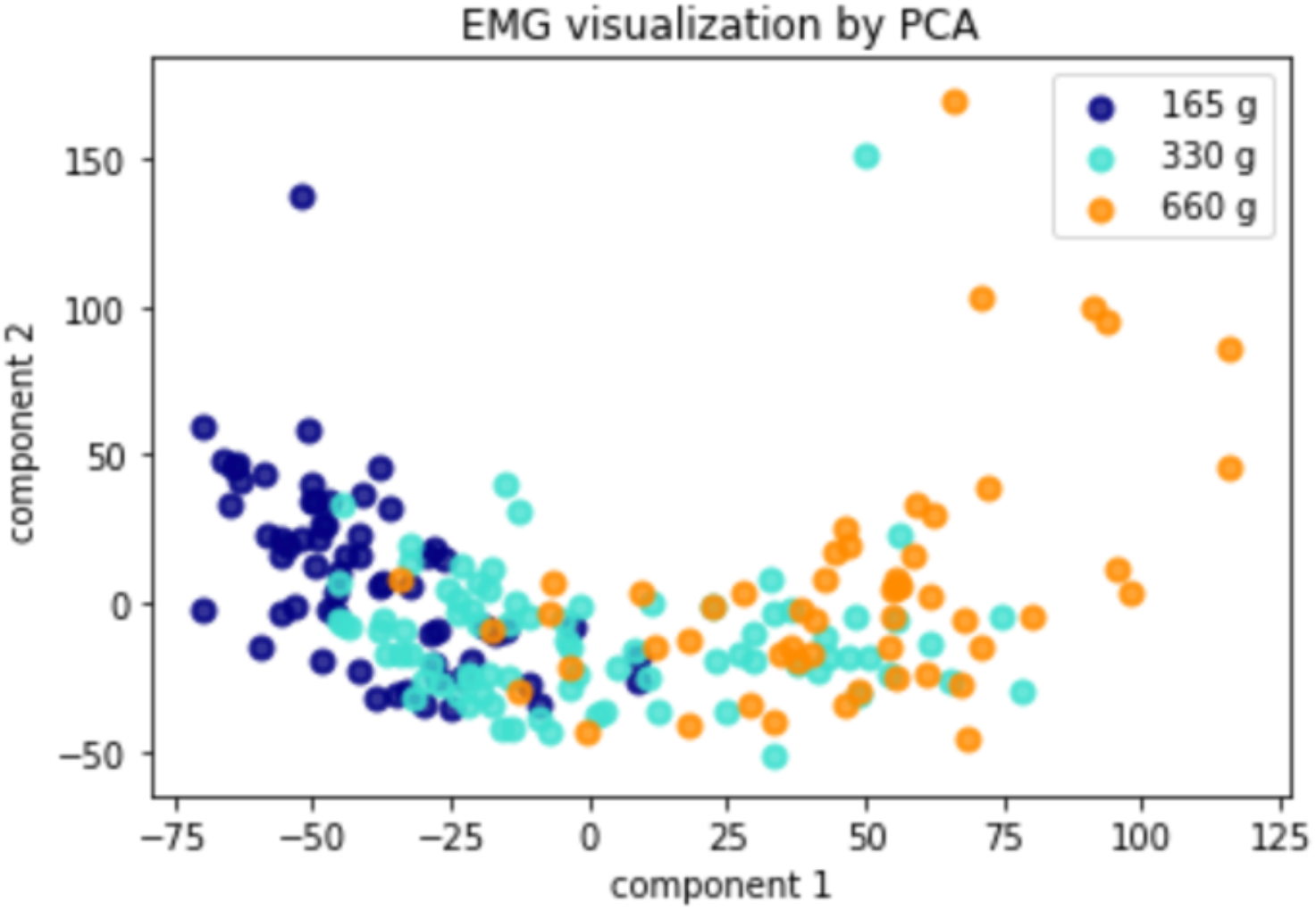
EMG visualization by two prominent components of PCA.

## 3. Results

This study utilized a PC based system and Python3 on a 3.4 GHz Intel^®^ CoreTM i7-6700 CPU. In this research, we first isolated 25% of the data as a separate test set. All the training as well as parameter and model selection was used only on the remaining 75%. Five-fold cross-validation was used to evaluate the classification accuracy of our model on the training dataset while performing a grid search over hyperparameters. Hyperparameters that yielded the best cross-validation accuracy were used to train a model over the entire training data. Finally, the trained model was fixed and then used to provide classification results over the test set that was isolated in the first step.

We report the accuracy of the classification using different muscles and segments, to test whether and to what extent our pipeline can classify different grasping tasks for the test data. We used 3 classifiers: Support vector machine SVM [21], k-Nearest Neighbors (k-NN) [28] and Linear Discriminant Analysis (LDA). The highest accuracy was obtained by linear SVM (Figure 5). We, therefore, used linear SVM for the remaining analysis. SVM with a linear kernel is a relatively simple classifier and offers good performance when the data is linearly separable. We also tested RBF SVM on our data, but the results were no better than linear SVM. So, we opted for the simpler linear SVM model.

**Figure 5:**
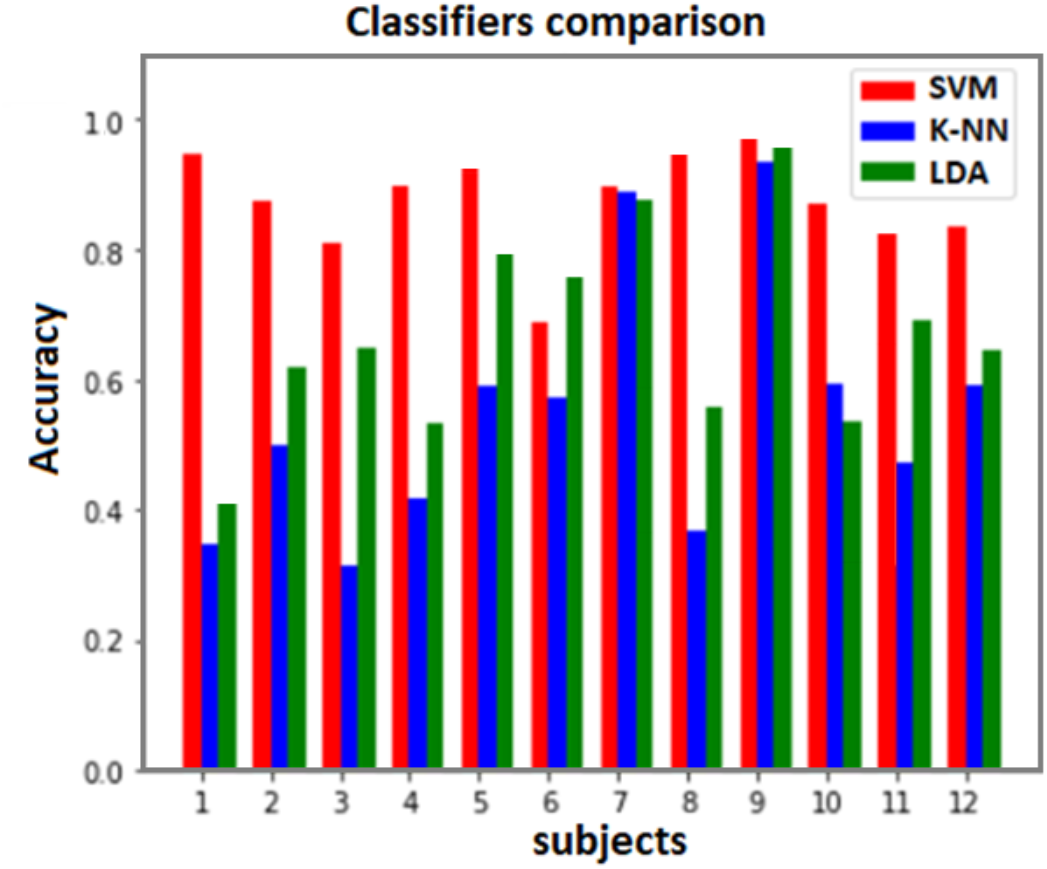
The accuracy camparison of different classifiers (SVM:red, k-NN:blue and LDA:greean) for all 12 subjects

Table 1 compares the accuracy of the classification of 3 objects with different weights using a single muscle and combination of all the 5 muscles with different segments length (i.e. (0-4s) 0 s: Touch event to 4 s after that). Based on the result, Brachioradialis yielded the highest classification accuracy and was thus the most informative muscle for identifying a specific weight. Combining all 5 muscles gives the best accuracy. We also focused on different intervals of selected segments to find the most important part of the grasping task. We found that the most informative of the selected segments are related to the first two seconds. This is when the subject starts lifting the weight and reaches the highest position. However, the value of 0.9 achieved by the combination of all muscles with the 4s segment.

Finally, we evaluated the output of the best classifier (SVM) on the data set using a confusion matrix and a receiver operating characteristic (ROC) curve [22, 23]. These plots provide a more global comprehensive view of the test. The diagonal elements of a confusion matrix represent the number of points for which the predicted label is equal to the true label, while off-diagonal elements are mislabeled by the classifier. The higher the diagonal values of the confusion matrix the better, indicating many correct predictions. We used the normalized confusion matrix because our classes are imbalanced (700 trials for 165g, 935 trials for 330, and 569 trials for 660 g). Figure (6)-a demonstrates that the best prediction for First Dorsal Interosseous muscle of Subject 1 is for class1(165 g). The ROC curve is a plot of the true-positive rate (y-axis) against the corresponding false positive rate (x-axis) on the independent test set. The ROC plot, therefore, describes test performance measured using sensitivity and 1-specificity at different thresholds. Figure (6)-b shows the ROC curves—i.e., the true-positive rate versus the false-positive rate for each class. The top-left corner of the plot is the ideal point—a false-positive rate of zero, and a true-positive rate of one. The area under the curve (AUC) of the ROC curve is often used to summarize test performance across all the threshold in the plot. A straight line connecting the extreme bottom-left (sensitivity (0,0)) and top-right (the chance diagonal (1,1)) describes a test with no discrimination (AUC=0.5, the same as flipping a coin). Based on this figure the AUC for 165 g is 0.99, 0.93 for 330 g and 0.97for 660 g. In this 3-class model, ROC for 165 g classified against 330 g and 660 g has the best result, which shows the slight change in higher weight is hard to detect rather than light weight.

**Figure 6:**
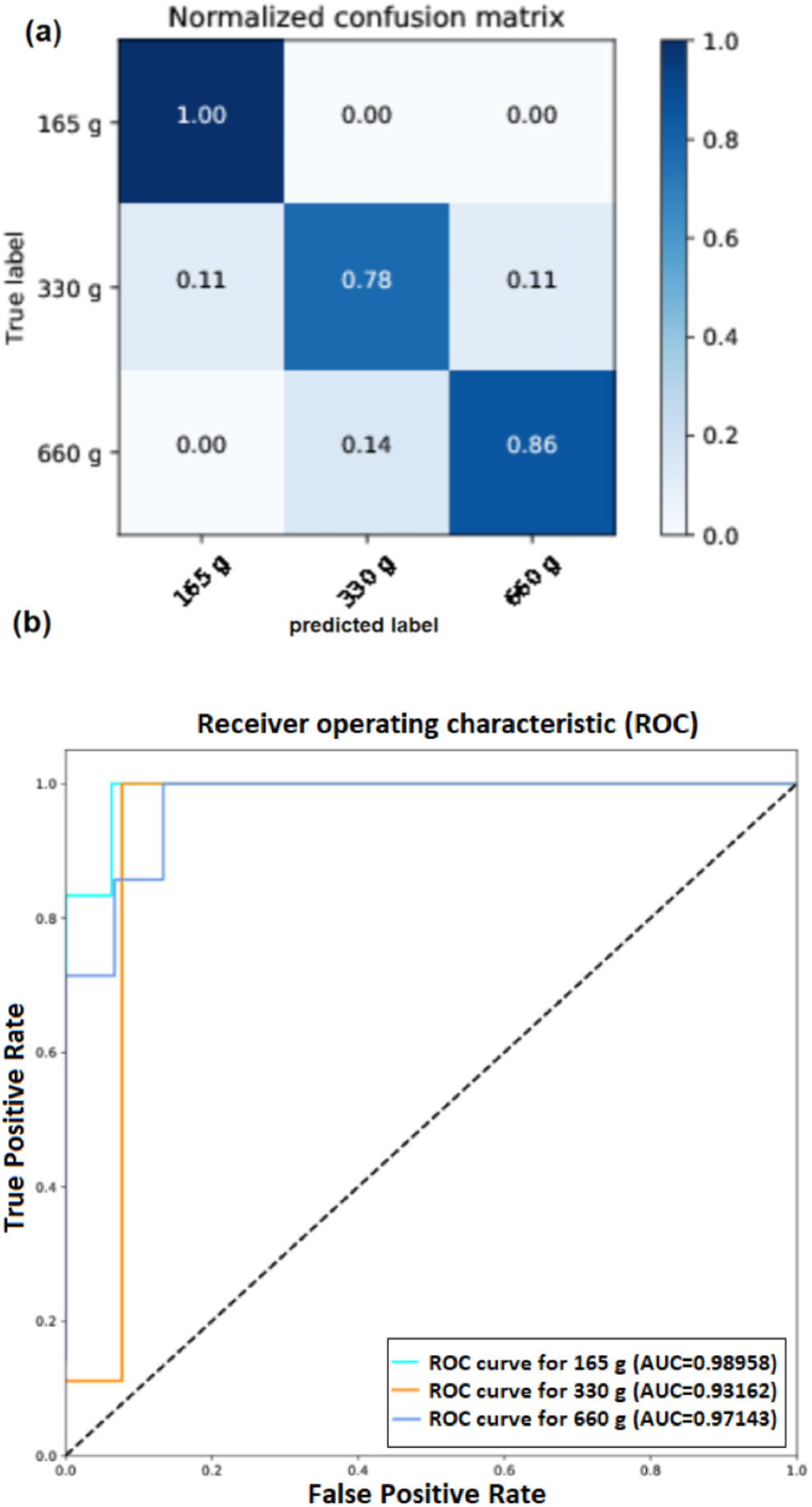
Evaluation of the proposed method for the classification of different weights (class 1: 165 g, class 2: 330 g, and class 3: 660g). (a) Normalized confusion matrix (b) Receiver operating characteristic (ROC) curve and Area Under the Curve (AUC)

## 4. Discussion and conclusion

This study proposes a pipeline for the pre-processing, feature selection, feature extraction and classification of different motor tasks (grasping three objects of different weights: 165, 330, and 660 g) based on EMG data alone. Grasping is an important topic in neuroscience and motor control, in biomechanics, and in robotics. We used a freely available dataset, which offers insights into human behavior as well as the control and design of artificial systems. Our pipeline achieves a classification accuracy of up to 90% on this 3-way classification. Feature selection is a very important stage in our procedure. We didn’t find informative data before movement onset, therefore, we locked on to the moment of touch and picked the window from this point to 4 seconds after that. We used the amplitude of EMG signals (time-domain features) for all 5 muscles. PCA was also important to extract the intrinsic data features. In our dataset 170 components were capable of describing 95% of the total variance. The results suggest that, during a grasp-and-lift task, the most informative single-muscle information was gained from the Brachioradialis, which seems plausible. Though the combination of all 5 muscles had the highest accuracy, pointing at additional information that exists in muscles other than the Brachioradialis (at least for linear SVM). Previous work on this dataset mainly attempted binary classification for the two a priori most distinct classes (165 and 660 g) by considering EEG and EMG signals and extracting frequency domain features with the same accuracy [14].

The methodology proposed in this paper can be easily adapted to each subject. It is also well suited to real-time operation, because the computational load in training and testing the model is relatively low. Our results might be of interest for motion and path planning, robotic learning and training, for rehabilitation, and for the investigation of human behavior. This methodology could also be a first step for a more-ambitious project of finding a mapping between EMG and EEG signals leading up to and during movement.

## 5. Ethical statement

We declare that this manuscript is original, has not published before and is not currently being considered for publication elsewhere. We know of no conflicts of interest associated with this publication, and has been no significant financial support for this work. As Corresponding Author, I confirm that the manuscript has been read and approved for submission by all the named authors.

## References

1. Kourtzi, Z., et al., Object-selective responses in the human motion area MT/MST. Nature neuroscience, 2002. 5(1): p. 17.

2. Chaffin, D.B., G. Andersson, and B.J. Martin, Occupational biomechanics. 1999: Wiley New York.

3. Grosse, P., M. Cassidy, and P. Brown, EEG–EMG, MEG–EMG and EMG–EMG frequency analysis: physiological principles and clinical applications. Clinical Neurophysiology, 2002. 113(10): p. 1523–1531.

4. Cifrek, M., et al., Surface EMG based muscle fatigue evaluation in biomechanics. Clinical Biomechanics, 2009. 24(4): p. 327–340.

5. Bashashati, A., et al., A survey of signal processing algorithms in brain–computer interfaces based on electrical brain signals. Journal of Neural engineering, 2007. 4(2): p. R32.

6. Pollard, N.S., et al. Adapting human motion for the control of a humanoid robot. in Robotics and Automation, 2002. Proceedings. ICRA’02. IEEE International Conference on. 2002. IEEE.

7. Lashgari, E. and E. Demircan, Electromyography Pattern Classification with Laplacian Eigenmaps in Human Running. World Academy of Science, Engineering and Technology, International Journal of Electrical, Computer, Energetic, Electronic and Communication Engineering, 2017. 11(4): p. 399–404.

8. Huang, Y., et al., Recent data sets on object manipulation: A survey. Big data, 2016. 4(4): p. 197–216.

9. Lorrain, T., N. Jiang, and D. Farina, Influence of the training set on the accuracy of surface EMG classification in dynamic contractions for the control of multifunction prostheses. Journal of neuroengineering and rehabilitation, 2011. 8(1): p. 25.

10. Alkan, A. and M. Günay, Identification of EMG signals using discriminant analysis and SVM classifier. Expert Systems with Applications, 2012. 39(1): p. 44–47.

11. Cisotto, G., et al., Classification of grasping tasks based on EEG-EMG coherence. arXiv preprint arXiv:1809.03300, 2018.

12. Canal, M.R., Comparison of wavelet and short time Fourier transform methods in the analysis of EMG signals. Journal of medical systems, 2010. 34(1): p. 91–94.

13. Murugappan, M., N. Ramachandran, and Y. Sazali, Classification of human emotion from EEG using discrete wavelet transform. Journal of Biomedical Science and Engineering, 2010. 3(04): p. 390.

14. Luciw, M.D., E. Jarocka, and B.B. Edin, Multi-channel EEG recordings during 3,936 grasp and lift trials with varying weight and friction. Scientific data, 2014. 1: p. 140047.

15. Phinyomark, A., C. Limsakul, and P. Phukpattaranont, A novel feature extraction for robust EMG pattern recognition. arXiv preprint arXiv:0912.3973, 2009.

16. Tkach, D., H. Huang, and T.A. Kuiken, Study of stability of time-domain features for electromyographic pattern recognition. Journal of neuroengineering and rehabilitation, 2010. 7(1): p. 21.

17. Englehart, K. and B. Hudgins, A robust, real-time control scheme for multifunction myoelectric control. IEEE transactions on biomedical engineering, 2003. 50(7): p. 848–854.

18. Wold, S., K. Esbensen, and P. Geladi, Principal component analysis. Chemometrics and intelligent laboratory systems, 1987. 2(1-3): p. 37–52.

19. Schölkopf, B., A. Smola, and K.-R. Müller. Kernel principal component analysis. in International Conference on Artificial Neural Networks. 1997. Springer.

20. Chu, J.-U., I. Moon, and M.-S. Mun, A real-time EMG pattern recognition system based on linear-nonlinear feature projection for a multifunction myoelectric hand. IEEE Transactions on biomedical engineering, 2006. 53(11): p. 2232–2239.

21. Kotsiantis, S.B., I. Zaharakis, and P. Pintelas, Supervised machine learning: A review of classification techniques. Emerging artificial intelligence applications in computer engineering, 2007. 160: p. 3–24.

22. Halligan, S., D.G. Altman, and S. Mallett, Disadvantages of using the area under the receiver operating characteristic curve to assess imaging tests: a discussion and proposal for an alternative approach. European radiology, 2015. 25(4): p. 932–939.

23. LeDell, E., M. Petersen, and M. van der Laan, Computationally efficient confidence intervals for cross-validated area under the ROC curve estimates. Electronic journal of statistics, 2015. 9(1): p. 1583.

